# Maf1 phosphorylation is regulated through the action of prefoldin-like Bud27 on PP4 phosphatase in *Saccharomyces cerevisiae*

**DOI:** 10.1101/2023.12.20.572514

**Authors:** F. Gutiérrez-Santiago, V. Martínez-Fernández, A.I Garrido-Godino, C. Colino-Palomino, A. Clemente-Blanco, C. Conesa, J. Acker, F. Navarro

**Author notes:** Laboratoire de Biologie Moléculaire et Cellulaire des Eucaryotes, UMR8226, CNRS, Institut de Biologie Physico-Chimique, Paris Sciences et Lettres Research University, Sorbonne Université, F-75005 Paris, France.

## Abstract

Bud27 is a prefoldin-like protein that participates in transcriptional regulation mediated by the three RNA polymerases in *Saccharomyces cerevisiae*. Lack of Bud27 significantly affects RNA pol III transcription, although the involved mechanisms have not been characterized. Here we show that Bud27 regulates the phosphorylation state of the RNA pol III transcriptional repressor, Maf1, influences its nuclear localization, and likely its activity. We demonstrate that Bud27 is associated with the Maf1 main phosphatase PP4 *in vivo,* and that this interaction is required for proper Maf1 dephosphorylation. Lack of Bud27 decreases the interaction among PP4 and Maf1, Maf1 dephosphorylation, and its nuclear entry. Our data uncover a new nuclear function of Bud27, identify PP4 as a novel Bud27 interactor and demonstrate the effect of this prefoldin-like on the posttranslational regulation of Maf1. Finally, our data reveal a broader effect of Bud27 on PP4 activity by influencing, at least, the phosphorylation of Rad53.

## INTRODUCTION

The prefoldin-like Bud27 and its orthologue URI are members of the prefoldin family of ATP-independent molecular chaperones, which function as scaffold proteins during the assembly of additional prefoldin family members (1,2). Bud27 and URI show nuclear and cytoplasmic localization (1,3,4), and associate with not only subunits Pfd2 and Pfd6 of the prefoldin complex, but also with Rpb5, a common subunit of the three RNA polymerases (1-3,5,6). Bud27 mediates the biogenesis and assembly of the three RNA pols in an Rpb5-dependent manner (3,7) and, similarly, URI participates in RNA pol II biogenesis (3,8-10). Bud27 and URI play multiple roles in transcription (2,5-7,11-14). However, only Bud27 has been reported to influence transcription by the three RNA pols (13,15), a process that impacts ribosome biogenesis, likely through a functional connection between Bud27 and the TOR pathway (15). Lack of Bud27 partially mimics the transcriptional response provoked by TOR pathway inactivation, (15) and Bud27 is proposed to impact some TOR-dependent processes, including cell wall integrity and autophagy induction (16). Both Bud27 and URI have been shown to participate in the TOR signaling cascade by coordinating nutrient availability with gene expression (13). More direct and specific roles of Bud27 in transcription have also been described. For example, Bud27 has been shown to be necessary in concert with the RSC chromatin remodeler complex for the elongation step of RNA pol II transcription (11). This functional relation between Bud27 and RSC also impacts chromatin organization and the polyadenylation site selection (17). The association between Bud27-RSC and RNA pol III has been proposed to influence its transcriptional activity (7) and lack of Bud27 has been shown to adversely affect the synthesis and probably the maturation of RNA pol III transcripts (7,15).

RNA pol III is a multiprotein complex that contains 17 subunits. It acts in concert with TFIIIB and TFIIIC, two basal transcription factors, for the synthesis of transcripts (18). Maf1, a protein conserved from yeast to humans, is the master negative regulator of RNA pol III-directed transcription in response to not only multiple stresses, but also to the inactivation of the TOR signaling pathway, and it interacts with distinct RNA pol III subunits and TFIIIB (19-23). Maf1 activity and cellular localization are regulated by its phosphorylation state. Under favourable growth conditions, Maf1 is mainly phosphorylated and located in the cytoplasm. Under repressive conditions, Maf1 dephosphorylation promotes its entry into the nucleus, where Maf1 binding to RNA pol III represses its transcriptional activity (20,24-26) and interferes with TFIIIB recruitment to promoters (27). Sch9 has been proposed as the major kinase for Maf1 phosphorylation, although PKA, CK2 and TORC1 have also been shown to mediate this process (24,28-32).

Protein Phosphatase 4 (PP4), a member of the phosphoprotein type 2A phosphatases family (PPPs), has been reported as the major phosphatase for Maf1 dephosphorylation(19). PP4 is an evolutionary conserved Ser/Thr phosphatase involved in several essential processes in eukaryotic cells, such as DNA repair (33,34). In *S. cerevisiae*, the PP4 complex comprises catalytic subunit Pph3, regulatory element Psy2 and regulatory/scaffold subunit Psy4 (35). In line with the role of PP4 in Maf1 dephosphorylation, the deletion of *PPH3* or *PSY2* genes impairs both Maf1 dephosphorylation and RNA pol III transcriptional repression after TOR pathway inhibition (36). Notably, PP4 associates with Maf1 *via* the interaction between Maf1 and Pph3 in both log-phase and stressed cells (36).

In this work, we investigated the functional relation between Bud27 and Maf1. We demonstrate that lack of Bud27 affects Maf1 dephosphorylation in response to stresses. We show that Bud27 interacts with PP4 phosphatase and that lack of Bud27 affects PP4-Maf1 interaction and Maf1 dephosphorylation, which alter Maf1 nuclear shuttling, and likely RNA pol III transcriptional repression. Our data also suggest a broader role of Bud27 in PP4 activity by affecting other PP4-dependent processes, such as DNA repair.

## MATERIALS AND METHODS

### Yeast strains and growth conditions

The *Saccharomyces cerevisiae* strains used in this study are listed in Table S1. Plasmids are listed in Table S2.

Media preparation, growth conditions, yeast transformation and genetic techniques were used as described elsewhere (37). Yeast strains were cultured at 30 °C in liquid YPD or synthetic SD medium supplemented with the specific nutrients required for auxotrophic deficiencies. Rapamycin treatment was performed by adding 400 nM of rapamycin to the cell cultures grown to exponential phase (OD_600_∼0.6) for 1 hour, unless otherwise specified.

Whenever indicated, cycloheximide (10 μg/ml) was added to cultures 10 minutes before rapamycin treatment to avoid Maf1 neo-synthesis due to stress (19). Other stresses were performed as follows: nutrient-deprived cells were grown in YPD until log phase and then washed 3 times with poor nutrient media (0.15X synthetic complete (SC), no glucose) and transferred to the same media for the indicated times, as previously described (36); dithiothreitol-treated cells were grown in YPD until log phase and then treated with 5 mM of DTT for the indicated times.

For the spotting assays, 10-fold serial dilutions of cultures were spotted onto YPD plates containing rapamycin, 4-Nitroquinoline 1-oxide (4-NQO), methyl methanesulfonate (MMS) or phleomycin at the indicated concentrations.

### Protein extraction, western blotting analyses and immunoreactive bands quantification

Whole cell protein extracts were prepared from 5 ml of the exponentially-grown cells (OD_600_ ∼0.7) using trichloroacetic acid and acid-washed glass beads as described (19). Protein extracts were analysed by SDS-PAGE and western blotting using different primary antibodies: anti-Maf1 (a gift from C. Conesa); anti-Pgk1 (459250, Invitrogen); PAP antibody (P1291, Merk); anti-c-Myc (9E10, Santa Cruz Biotechnology or C3956, Merk); anti-HA (3F10, Roche); anti-Rad53 (ab104232, Abcam). To facilitate Psy4, Psy2 and Pph3 phospho-bands separation, a final concentration of 10 μM of Phos-Tag (Wako, 300-93523) was added to the gel. Immunoreactive bands were revealed using SuperSignal® West Femto (Thermo Scientific, 10391544), followed by exposure to ECL Hyperfilm (GE Healthcare, 28-9068-37) and then quantified by densitometry using the Image Studio LITE software from the images acquired with an office scanner.

### Protein immunoprecipitation and TAP purification

For immunoprecipitation purposes, the whole-cell extracts were prepared from 200 ml of exponentially-grown cells (OD_600_∼0.6) as described (38). Purifications were carried out from 150 μl of the whole-cell extract (2 mg protein) per experiment using 37 µl of Dynabeads M-280 sheep anti-rabbit IgG (Invitrogen) and 2 µl of the anti-LytA antibody (Biomedal). For TAP purification, the same protocol was followed with 50 µl of Dynabeads Pan Mouse IgG (Invitrogen). The affinity-purified proteins were released from beads by boiling for 10 min and were analysed by western blotting. The intensities of the immunoreactive bands were quantified by densitometry using the software TOTALLAB from the images acquired with an office scanner.

### Chromatin-enriched fractions and western blotting analyses

The yChEFs procedure was followed to prepare chromatin-enriched fractions (39,40) using 50 ml of the exponentially-grown cultures in YPD medium with or without rapamycin addition for 1 h (OD600∼0.6–0.8). The final chromatin-bound proteins were resuspended in 1X SDS-PAGE sample buffer, boiled and analysed by western blotting with different antibodies: anti-c-Myc (9E10, Santa Cruz Biotechnology or C3956, Merk); anti-Maf1 (a gift from C. Conesa); anti-Pgk1 (459250, Invitrogen); anti-H3 (ab1791; Abcam).

The Image Studio LITE software was utilised to quantify the intensities of the immunoreactive bands.

### Immunolocalization and fluorescence microscopy

Immunolocalization and fluorescence microscopy were performed essentially as previously described (41) from cells grown at 30°C in YPD until log phase (OD_600_∼0.6) using 1:100 dilutions of primary antibodies anti-c-Myc (9E10, Santa Cruz Biotechnology), anti-HA (3F10, Roche), polyclonal or anti-TAP (Thermo Fisher), followed by incubation with a 1:100 dilution of the secondary antibody Alexa Fluor^TM^ 488 (Thermo Fisher). Slides were washed 3 times with PBS and incubated for 5 min with 50 µl of 1 mg/ml DAPI (in PBS), washed again and finally covered with Vectashield (Vector Laboratories) mounting solution. Fluorescence intensity was scored with a fluorescence microscope (Olympus BX51).

### RNA extraction, reverse transcription, qPCR analysis and northern blotting

Total RNA was extracted from 50 ml of cell cultures grown in YPD until log phase (OD_600_∼0.6) following the hot phenol acid extraction protocol as described (15). First-strand cDNA was synthesized from 500 ng of total RNA with the iScript cDNA synthesis kit (Bio-Rad) in accordance with the manufacturer’s instructions, albeit with some modifications. The reaction mixtures containing 500 ng of total RNA, 4 μl of the iScript reaction mix 5X, 500 ng of total RNA and 10 pmol of the specific reverse primer to amplify the gene of interest were heated at 65°C for 10 minutes. Tubes were kept on ice for 5 minutes before adding iScript reverse transcriptase. Each sample was subjected to the same reaction without reverse transcriptase to be employed as a negative control for genomic DNA contamination.

Real-time qPCR was performed in a CFX-384 Real-Time PCR instrument (BioRad) with the ‘SsoFast^TM^ EvaGreen ® Supermix’ EvaGreen detection system (Bio-Rad). Reactions were performed in a 5 µl total volume at the 1:100 dilution of synthesized cDNA. Each sample was analysed in triplicate with at least three independent biological replicates per sample. Values were normalized in relation to the steady-state levels of the 18S cDNA levels. The used oligonucleotides are listed in Supplemental Table S3.

For northern blotting, equal amounts of total RNA (5 μg) were loaded on 7% polyacrylamide-8M urea gels. Specific oligonucleotides were 5′-end labeled with [γ-32P] ATP and used as probes. The sequences of the oligonucleotides used for northern hybridization are listed in Supplemental Table S4. The Phosphorimager analysis was performed in a FLA-5100 imaging system (Fujifilm) at the Biology Service (CITIUS) of the University of Seville (Spain).

## RESULTS

### Maf1 dephosphorylation under stress conditions depends on Bud27

Bud27 inactivation provokes a transcriptional response that partially mimics TOR pathway inactivation (15), a situation that leads to RNA pol III transcription repression *via* Maf1 (20). As all these elements are functionally related to the TOR pathway, we wondered whether Bud27 could influence Maf1 regulation.

As Maf1 is dephosphorylated and represses RNA pol III (36) under unfavourable growth conditions and stress, we investigated the dephosphorylation kinetics of Maf1 in wild-type or *bud27Δ* cells upon rapamycin addition for up to 120 min to repress the TOR pathway. As expected (36,42), both phosphorylated and dephosphorylated Maf1 were detected by western blotting in the extracts prepared from exponentially-grown cells. However, the ratios of the two Maf1 forms were not similar in each strain (Figure 1A). In the *bud27Δ* cells, the steady-state levels of phosphorylated Maf1 were higher than the dephosphorylated form unlike the wild-type cells. Whereas Maf1 dephosphorylation occurred rapidly as expected in the wild-type strain, Maf1 dephosphorylation slowed down when Bud27 was absent, and even seemed incomplete even after 2 h of rapamycin treatment. The accumulation of phosphorylated Maf1 under normal growth conditions, as well as the defect in dephosphorylation upon stress in the *bud27Δ* cells, suggest that Bud27 may interfere with Maf1 dephosphorylation.

**Figure 1.**
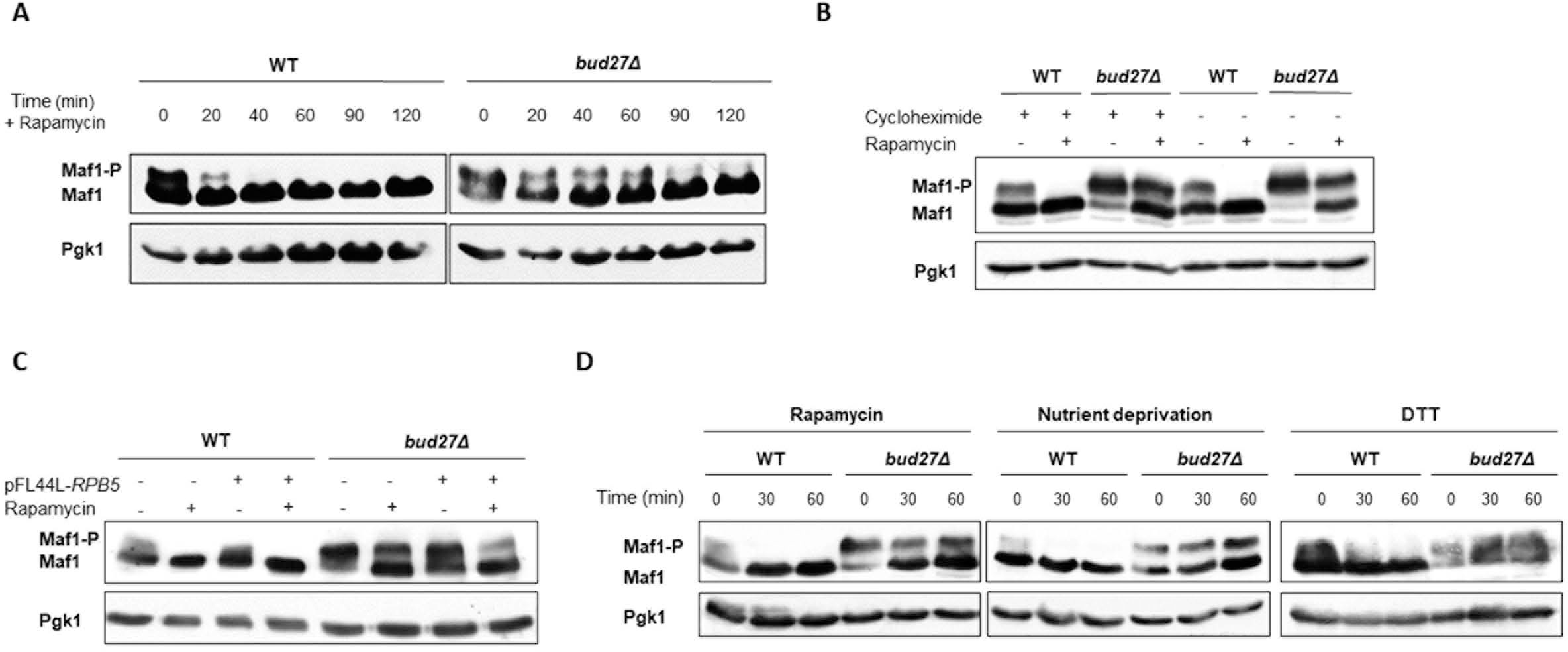
Maf1 dephosphorylation upon TOR pathway inactivation depends on Bud27. A) Analysis of Maf1 dephosphorylation in the *bud27Δ* cells and their isogenic wild-type strain after TOR pathway inactivation by rapamycin. The whole-cell extracts prepared from the wild-type (WT) or *bud27Δ* cells grown to the log phase (time 0) or after rapamycin treatment for the indicated times were analysed by western blotting with specific antibodies against Maf1 or Pgk1 used as an internal control. B) Effects of cycloheximide on Maf1 dephosphorylation. Western blotting analysis of Maf1 as described in panel A, but with the addition of cycloheximide to avoid Maf1 neo-synthesis 10 min before rapamycin treatment for 1 hour. C) Analysis of Maf1 dephosphorylation upon *RPB5* overexpression. Western blotting analysis of Maf1 after rapamycin treatment for 1 h in cells transformed with a multicopy plasmid (pFL44) carrying *RPB5* or not, as indicated. D) Analysis of Maf1 dephosphorylation under different stress conditions. Western blotting analysis of Maf1 after rapamycin treatment, a shift to SC medium without glucose (nutrient deprivation) or DTT addition for the indicated times.

To rule out that larger amounts of phosphorylated Maf1 did not result from the newly translated protein, cells were pretreated briefly with cycloheximide which blocks translation and does not cause Maf1 dephosphorylation (19,36). Once again, accumulation of phosphorylated Maf1 under normal growth conditions and a defect in dephosphorylation upon stress were observed in the *bud27Δ* cells (Figure 1B). These results demonstrated that the defect in Maf1 dephosphorylation detected in the *bud27Δ* cells did not depend on Maf1 neosynthesis.

Lack of Bud27 affects the correct assembly and the nuclear localization of the three RNA pols, and these defects are offset by the overexpression of *RPB5*, the gene that codes for the Rpb5 common subunit of RNA pols (3). As shown in Figure 1C, the defect in Maf1 dephosphorylation was independent of RNA pol III assembly because it was still observed in the *bud27Δ* cells when *RPB5* was overexpressed.

Finally, we wondered whether the Maf1 dephosphorylation defect observed in the *bud27Δ* cells was either specific to rapamycin treatment or could be extended to other stresses. Maf1 phosphorylation status was analysed in the wild-type or *bud27Δ* cells grown under nutrient deprivation conditions (lack of glucose) or with endoplasmic reticulum stress (treatment with dithiothreitol, DTT) because these treatments have been reported to cause Maf1 dephosphorylation and RNA pol III repression (36,43). As shown in Figure 1D, both stress conditions resulted in a similar Maf1 dephosphorylation defect to that observed upon the rapamycin treatment of the *bud27Δ* cells.

Taken together, these data indicate that optimal Maf1 dephosphorylation depends on Bud27.

### Maf1 nuclear entry depends on Bud27

Under rapamycin addition and TOR pathway repression, Maf1 dephosphorylation is accompanied by its accumulation in the nuclear compartment (19,36).We wondered whether the dephosphorylation defect observed when Bud27 was absent could lead to Maf1 mislocalization because Maf1 dephosphorylation is a prerequisite for its translocation to the nucleus (19,36).

Thus we investigated Maf1 cellular localization after rapamycin addition in the wild-type and *bud27Δ* cells containing a functional Maf1-GFP tagged version of the protein. As shown in Figure 2A, Maf1 accumulation in the nucleus of the wild-type cells occurred within 30 min after TOR pathway repression. When Bud27 was lacking, Maf1-GFP remained mainly cytoplasmic, even after 90 min of rapamycin treatment, which suggests that Maf1-GFP nuclear translocation slowed down. The western blotting analysis of the same cell extracts (Figure 2B) in the absence of Bud27 indicated that Maf1-GFP dephosphorylation slowed down, as observed for endogenous Maf1 (Figure 1A). These data suggest that Maf1 dephosphorylation could be a prerequisite for its nuclear localization, which falls in line with previous published results (24,36) and indicate that Maf1 nuclear accumulation depends on Bud27.

**Figure 2.**
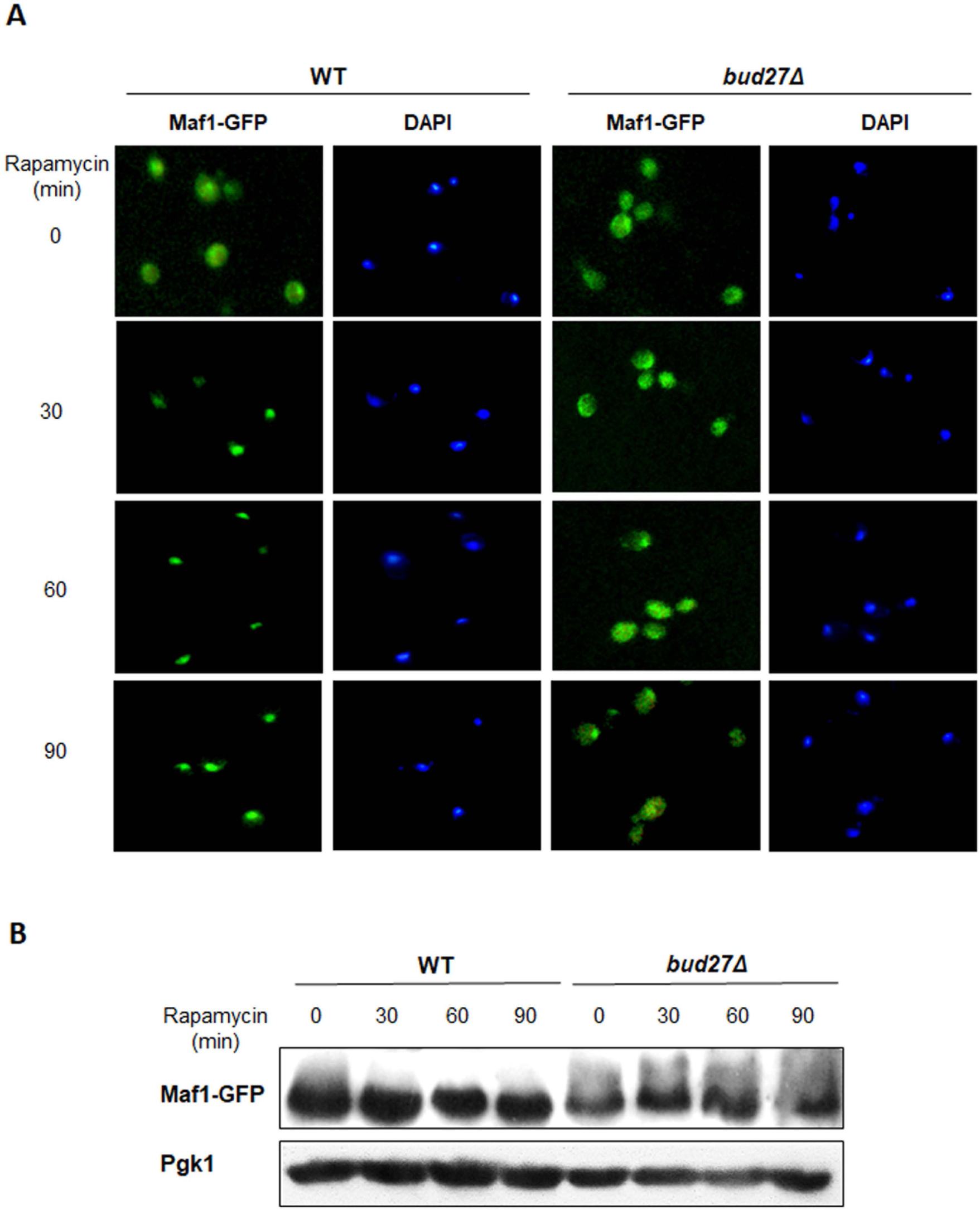
Maf1 nuclear entry and accumulation depends on Bud27. A) Live-cell imaging of Maf1-GFP in the wild-type or *bud27Δ* cells grown to the log phase (time 0) or treated with rapamycin for the indicated times. B) Analysis of Maf1-GFP dephosphorylation. The protein samples from the experiment in panel A were analysed by western blotting with the anti-Maf1 and anti-Pgk1 antibodies.

### Functional and genetic analyses suggest a functional relation between Bud27 and phosphatase PP4

We next investigated whether the known Maf1 kinases or phosphatases (19,20,24,26,32,36) could be involved in the dephosphorylation defect observed in the absence of Bud27.

To this end, we analysed the Maf1 dephosphorylation profile under TOR pathway repression by rapamycin addition in the wild-type and *bud27Δ* cells, as well as in the single mutants of several phosphatases (PP4 and PP2A) or kinases (PKA, Sch9, TORC1 and CK2), or in combination with *bud27Δ*. As shown in Figure 3A, a *pph3Δ* mutant (PP4 phosphatase) showed a similar alteration in Maf1 dephosphorylation as the *bud27Δ* cells and no additional defect was observed in the double mutant *pph3Δ bud27Δ*, which implies that both proteins were involved in the same dephosphorylation pathway of Maf1. Furthermore, no significant alterations were detected for PP2A phosphatase. These data agree with those previously reported for a *pph3Δ* mutant (36) and point out that PP4 is the major phosphatase involved in Maf1 dephosphorylation. In addition, no major defects in Maf1 dephosphorylation were observed in the PKA, TORC1 or CK2 mutant strains, unlike the strain deleted for *SCH9*, the proposed major kinase of Maf1 (32). Although no clear phosphorylated Maf1 was detected in the single *sch9Δ* mutant, phosphorylated Maf1 was observed in the *sch9Δ bud27Δ* double mutant, which suggests the existence of other kinases involved in the phosphorylation of this protein, which could be CK2 or PKA as previously proposed (26,30).

**Figure 3.**
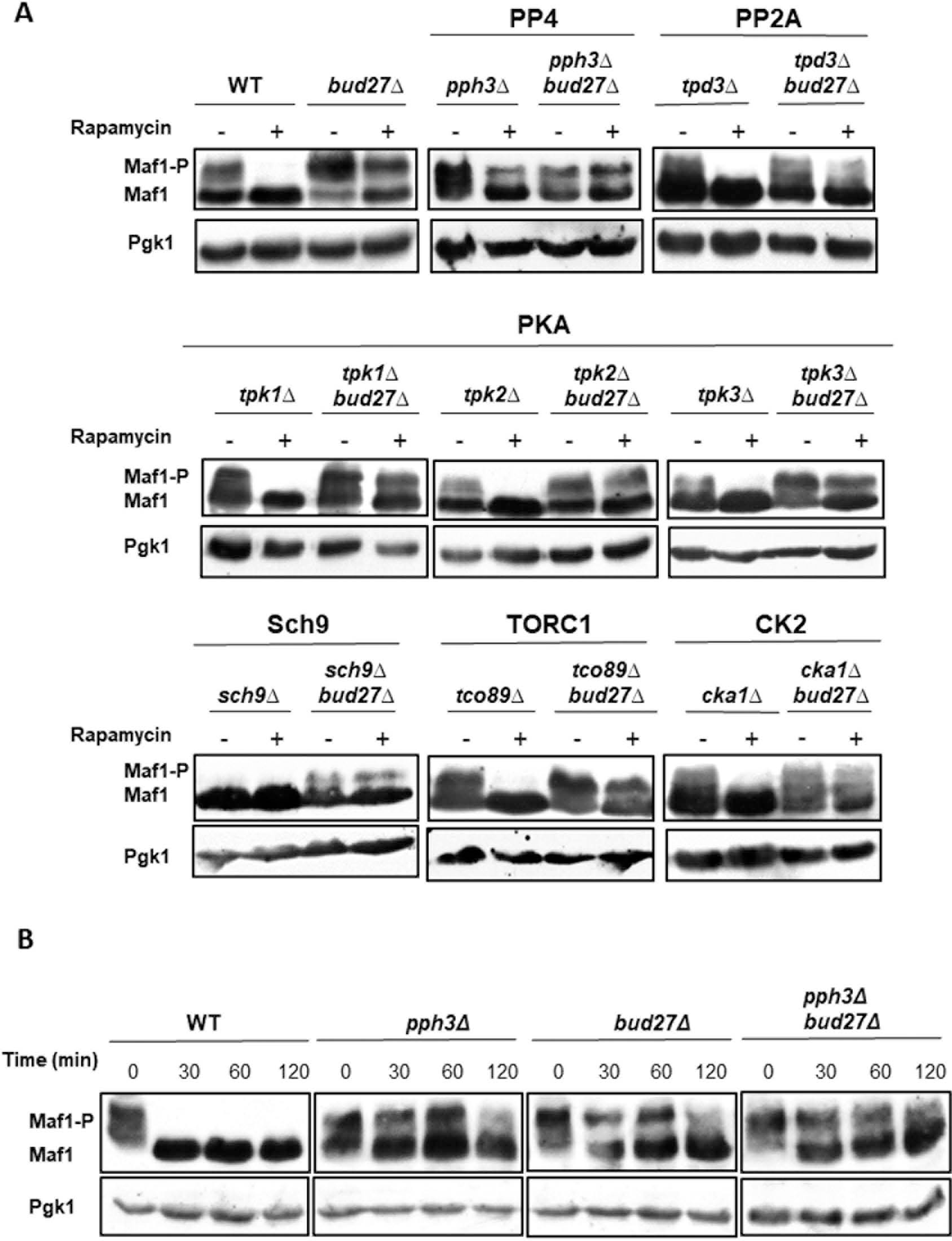
Bud27 and Pph3 are functionally related. A) Analysis of Maf1 dephosphorylation in the mutant strains of the Maf1-related phosphatases (PP4, PP2A) or kinases (PKA, Sch9, TORC1 or CK2) in combination, or not, with *bud27Δ*. Western blot analysis of Maf1 after rapamycin treatment for 1 h in the indicated mutants. B) Analysis of Maf1 dephosphorylation by western blotting in the wild-type (WT), *pph3Δ, bud27Δ* and *pph3Δ bud27Δ* cells after rapamycin treatment for the indicated times. The anti-Maf1 and anti-Pgk1 antibodies were used.

To investigate whether Bud27 and Pph3 act in concert or in different Maf1 dephosphorylation steps, we analysed the dephosphorylation kinetics of Maf1 in the *bud27Δ*, *pph3Δ* and *pph3Δ bud27Δ* mutant cells, and also in the isogenic wild-type strain. The results in Figure 3B revealed a similar Maf1 dephosphorylation pattern in both the single and double mutants, which implies that Bud27 and Pph3 may act in the same Maf1 dephosphorylation pathway.

Moreover, and in line with Bud27 participating in the TOR pathway and Maf1 dephosphorylation, positive genetic interactions were noted with the mutants of PP4 or PP2A phosphatases when cells were grown in the presence of rapamycin (Figure 4). On the contrary, negative genetic interactions were observed between the *bud27Δ* mutant and the mutants of the Sch9, CK2 and TORC1 kinases, whereas no differences were detected for PKA kinase (Figure 4). These results agree with the previously reported role of PP4/PP2A and Sch9/CK2 in Maf1 phosphorylation regulation (19,26,32,36).

**Figure 4.**
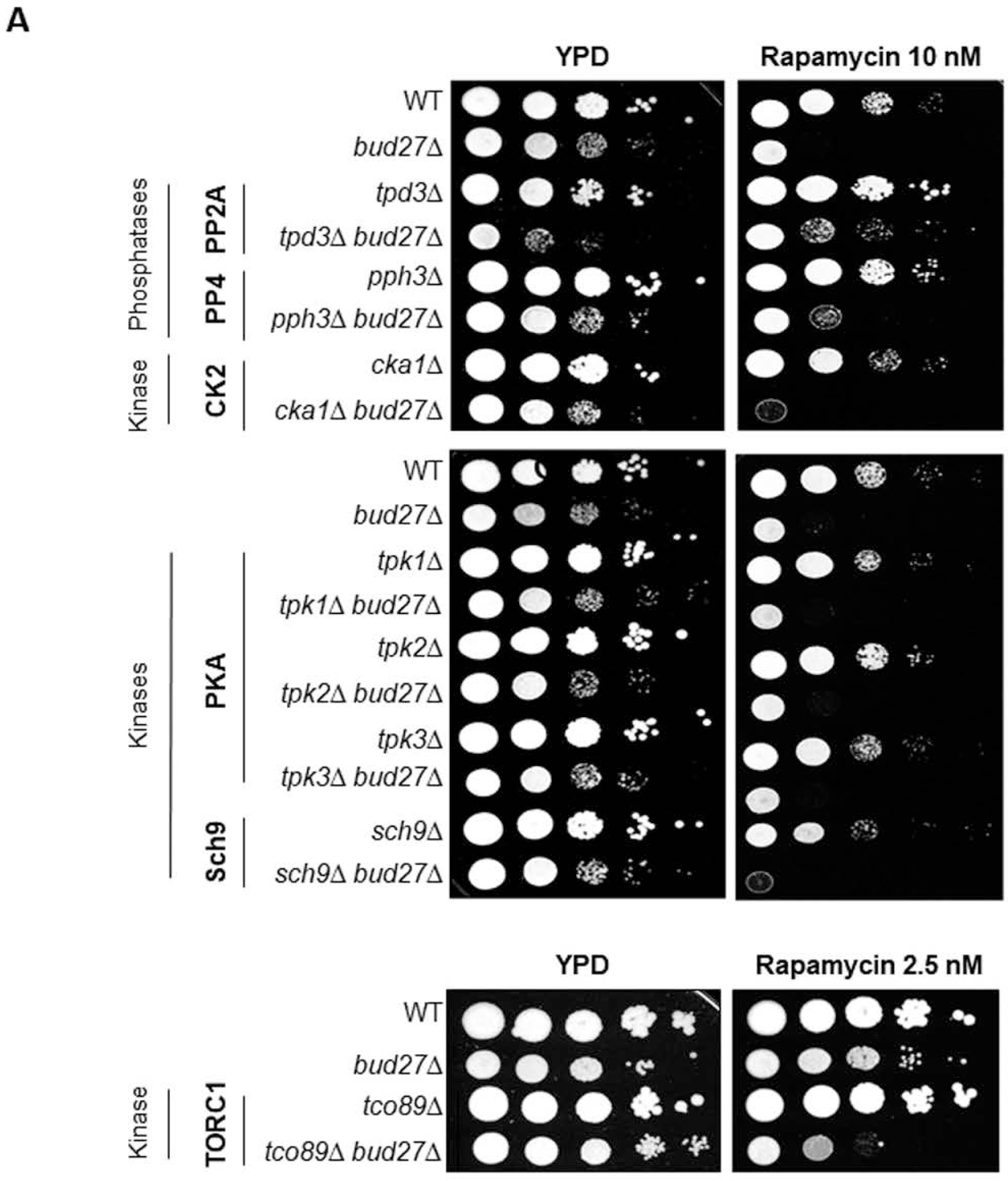
Genetic interactions between *BUD27* and the Maf1-related phosphatases or kinases. Genetic interactions between *BUD27* and *TPD3* (PP2A), *PPH3* (PP4), *CKA1* (CK2), *TPK1, TPK2* and *TPK3* (PKA), *SCH9* or *TCO89* (TORC1) were monitored by spotting assays. Serial dilutions of the indicated cell cultures were spotted onto YDP plates containing, or not, rapamycin at the indicated concentrations. Plates were incubated at 30°C.

### Bud27 associates with PP4 phosphatase *in vivo*

To explore the mechanisms that govern the action of Bud27 and PP4 on Maf1 phosphorylation and nuclear entry, we investigated the association between PP4 and Bud27.

We analysed whether Bud27 and PP4 were associated *in vivo*. We performed co-immunoprecipitation (co-IP) analyses on extracts prepared from the cells expressing both an ectopic TAP-tagged Pph3 (Pph3-TAP) from a pTet-off promoter and an endogenous tagged Bud27-LytA, grown either with or without rapamycin addition for 1 h. As shown in Figure 5A, Bud27 co-purified with Pph3-TAP after the immunoprecipitation with Dynabeads Pan Mouse IgG. We also corroborated these results by LytA immunoprecipitation from the cells expressing both endogenous tagged Bud27-LytA and Pph3-Myc proteins (Figure 5B). In both experiments, interactions were not altered when the TOR cascade was inhibited by rapamycin addition.

**Figure 5.**
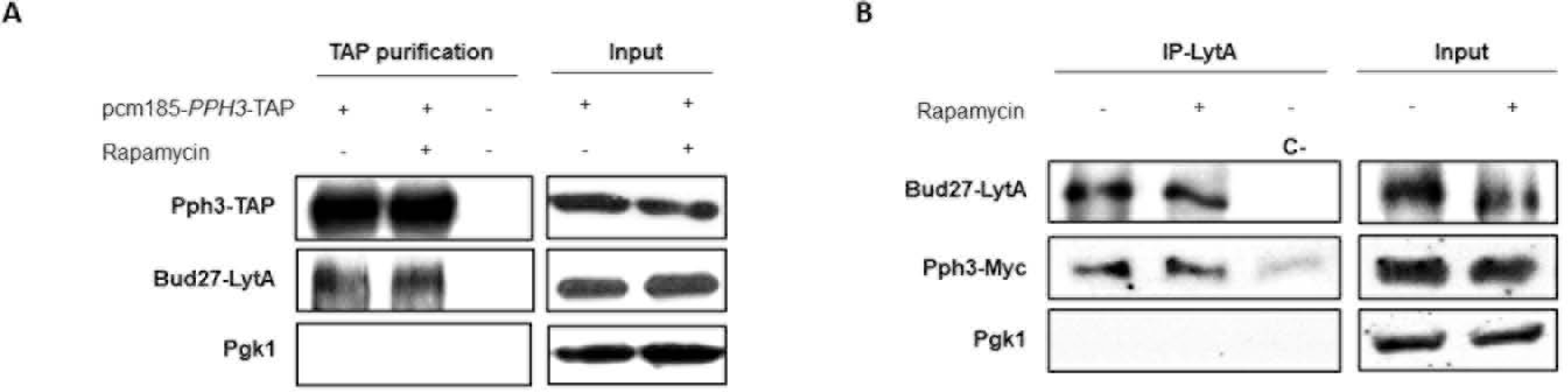
Bud27 interacts with Pph3 and mediates PP4-Maf1 association. A) The wild-type cells containing a LytA-tagged version of Bud27 and a TAP-tagged version of Pph3 (expressed from the pCM185-*PPH3*-TAP plasmid) were grown to the log phase and then treated (or not) with rapamycin for 1 hour. Whole-cell extracts (Input) were subjected to pull-down with Dynabeads Pan-Mouse. The obtained immunoprecipitates (TAP purification) were analysed by western blotting, performed with antibodies against TAP, LytA or Pgk1 as an internal control. B) The whole-cell extracts (Input) from the wild-type (WT) cells containing a LytA-tagged version of Bud27 and a Myc-tagged version of Pph3 treated (or not) with rapamycin for 1 hour were subjected to pull-down with anti-LytA antibodies. The obtained immunoprecipitates (IP-LytA) were analysed by western blotting, performed with antibodies against LytA, Myc or Pgk1. A LytA-untagged strain (C-) was used as a control.

These data strongly indicate that Bud27 and Pph3 are associated *in vivo*. However, as Pph3 is a component of the PP4 phosphatase complex, we could not conclude whether the interaction between Bud27 and Pph3 was direct.

### The association between PP4 phosphatase and Maf1 depends on Bud27 and influences the level of chromatin-associated Maf1 and RNA pol III transcription repression

We next wondered whether Bud27 could interfere with the interaction between Pph3 and Maf1 that has been reported previously (36) using co-immunoprecipitation experiments with cells overexpressing both the Pph3-TAP protein and Maf1. To do so, we performed similar experiments with the wild-type or *bud27Δ* mutant cells. We confirmed a physical interaction between Pph3 and Maf1 *in vivo,* but it showed a ≈70% decrease in their association in the absence of Bud27 (Figure 6A). Furthermore, inhibition of the TOR cascade by rapamycin addition had no effect on the Maf1/Pph3 interaction involving both strains (Figure 6).

**Figure 6.**
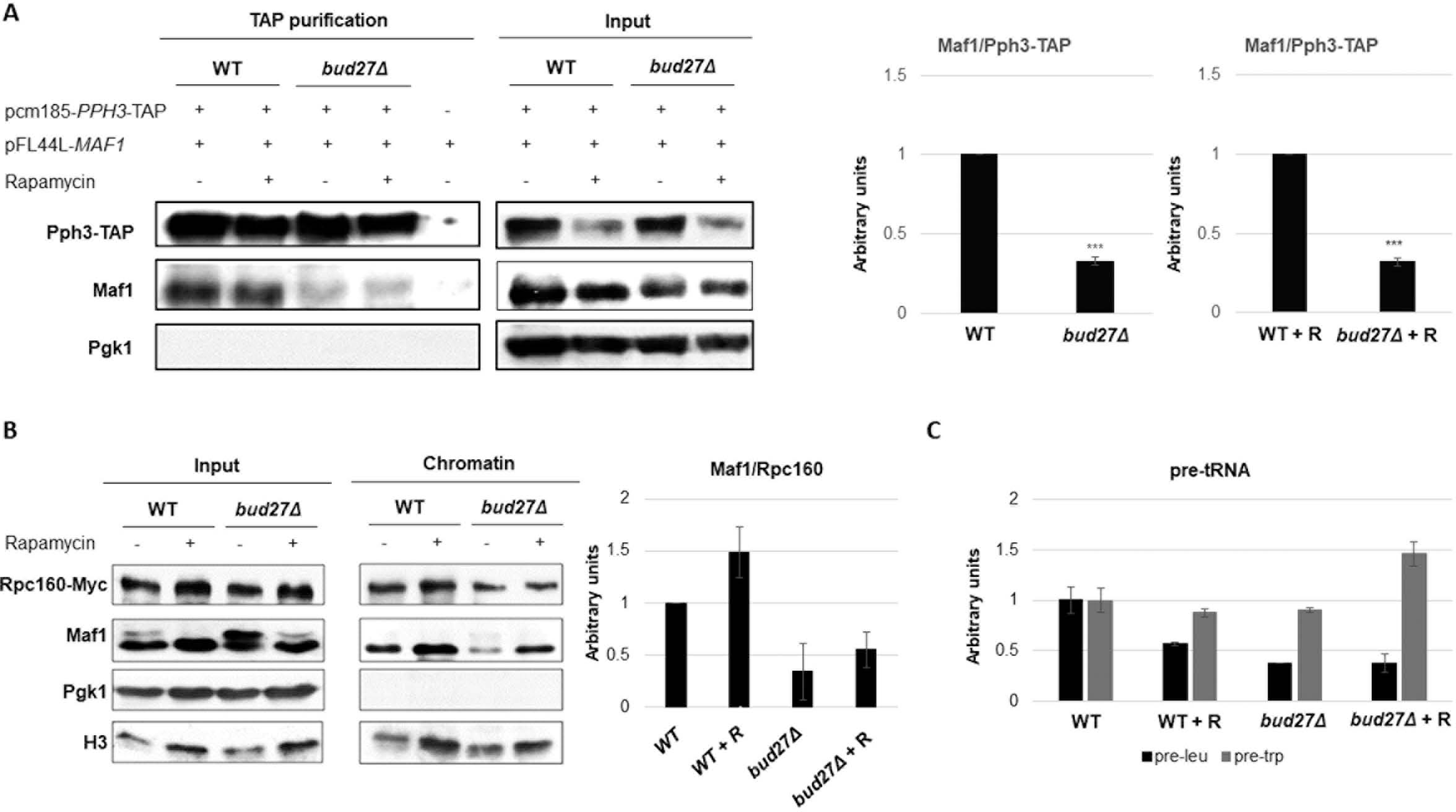
Lack of Bud27 affects PP4-Maf1 association and RNA pol III repression. A) Left panel. The whole cell extracts (Input) from the wild-type (WT) or *bud27Δ* cells carrying a plasmid for Maf1 overexpression (pFL44L-*MAF1*) and a pCM185-*PPH3*-TAP plasmid (or not as indicated) and treated (or not) with rapamycin for 1 hour were subjected to pull-down with Dynabeads Pan-Mouse. The obtained immunoprecipitates (TAP purification) were analysed by western blotting with antibodies against TAP, Maf1 or Pgk1. Right panel. Intensities of the immunoreactive bands from the western blotting shown in the left panel were quantified by densitometry. Graphs represent the median and standard deviation of three independent biological replicates. Statistical significance by t-Student, (∗∗∗) P < 0.0001. +R: + Rapamycin treatment. B) Left panel: chromatin-enriched fractions were obtained by the yChEFs procedure (39,40) from the wild-type or *bud27Δ* cells expressing Rpc160-Myc grown to the log phase and then treated (or not) with rapamycin for 1 hour. The proteins from chromatin and the whole cell-extracts (Input) were analysed by western blotting with antibodies against Myc, Maf1 and Histone H3 or Pgk1 as a control of chromatin or cytoplasm, respectively. Right panel: intensities of the immunoreactive bands from the western blottings shown in left panel were quantified by densitometry. Graphs represent the median and standard deviation of three independent biological replicates. C) RT-qPCR to analyse the neosynthesis of tRNALeu3 (pre-leu) and tRNATrp (pre-trp) from the wild-type or *bud27Δ* cells grown to the log phase and then treated (or not) with rapamycin for 1 hour. rRNA 18S was used as a normalizer in RT-qPCR. Experiments corresponded to three independent biological replicates. Bars represent standard deviation.

Taken together, our results indicate that the Bud27-PP4 phosphatase interaction is required for an optimal Pph3/Maf1 association, and suggest that the Maf1 dephosphorylation defect detected in the absence of Bud27 is probably due to the effect of Bud27 on PP4.

According to these data, we hypothesized that the defect in the Maf1-PP4 association provoked by lack of Bud27 could alter the Maf1 level associated with chromatin (*via* RNA pol III) and the transcriptional repression that occurs by inhibiting the TOR cascade. By western blotting, we first analysed the amount of Maf1 associated with chromatin in relation to the amount of the Rpc160 (Rpc160-Myc) subunit of RNA Pol III in the wild-type and *bud27Δ* mutant cells (with or without rapamycin addition for 1 h). This analysis was conducted with the chromatin-enriched fractions obtained by following the yChEFs procedure (39,40). As depicted in Figure 6B, and as expected, the Maf1/Rpc160 ratio increased upon TOR cascade inhibition in the wild-type cells. Furthermore, although the Maf1/Rpc160 ratio also increased with TOR cascade inhibition in the *bud27Δ* mutant cells, it was lower than that in the wild-type cells. In addition, similar observations were made under the non repressing conditions. These data correlated with the decrease in the nuclear amount of Maf1 found in the *bud27Δ* mutant cell, as shown in Figure 2.

Based on our above results, we speculated that RNA pol III transcription repression must be affected. To investigate the consequences of lack of Bud27 on RNA pol III repression, we isolated RNA from the wild-type and *bud27Δ* mutant cells treated or not with rapamycin for 1 h. Then we performed an RT-qPCR analysis for two pre-tRNA species: pre-tRNA-Leu3 and pre-tRNA-Trp. Figure 6C shows that repression of the pre-tRNAs neosynthesis under rapamycin addition occurred as expected in the wild-type strain. Furthermore these data revealed that *bud27Δ* was defective for RNA pol III transcriptional repression. Similar results were observed with the northern blotting experiments (Figure S1).

Taken together, these results are consistent with a model in which lack of Bud27 affects Maf1 dephosphorylation *via* PP4 phosphatase and alters proper RNA pol III transcriptional repression.

### Lack of Bud27 does not affect the composition or stoichiometry of the PP4 complex

Bud27 has been proposed to act as a co-chaperone in several processes (1,44). As we showed that Bud27 is associated with PP4 phosphatase (Figures 5A and 5B), we next explored whether lack of Bud27 could alter the composition, stoichiometry or stability of the PP4 complex. PP4 phosphatase was immunoprecipitated from the wild-type or *bud27Δ* cells expressing the Pph3-HA, Psy2-Myc and Psy4-TAP tagged subunits of PP4. As shown in Figure 7A, no significant differences in the amounts of the PP4 subunits were observed in either the cell crude extracts prepared from the wild-type or *bud27Δ* strains grown at 30°C (Figure 7A, Input) or the different immunoprecipitations (Figure 6A, 30°C). All this indicates that the composition or subunit stoichiometry of PP4 was not altered in the absence of Bud27.

**Figure 7.**
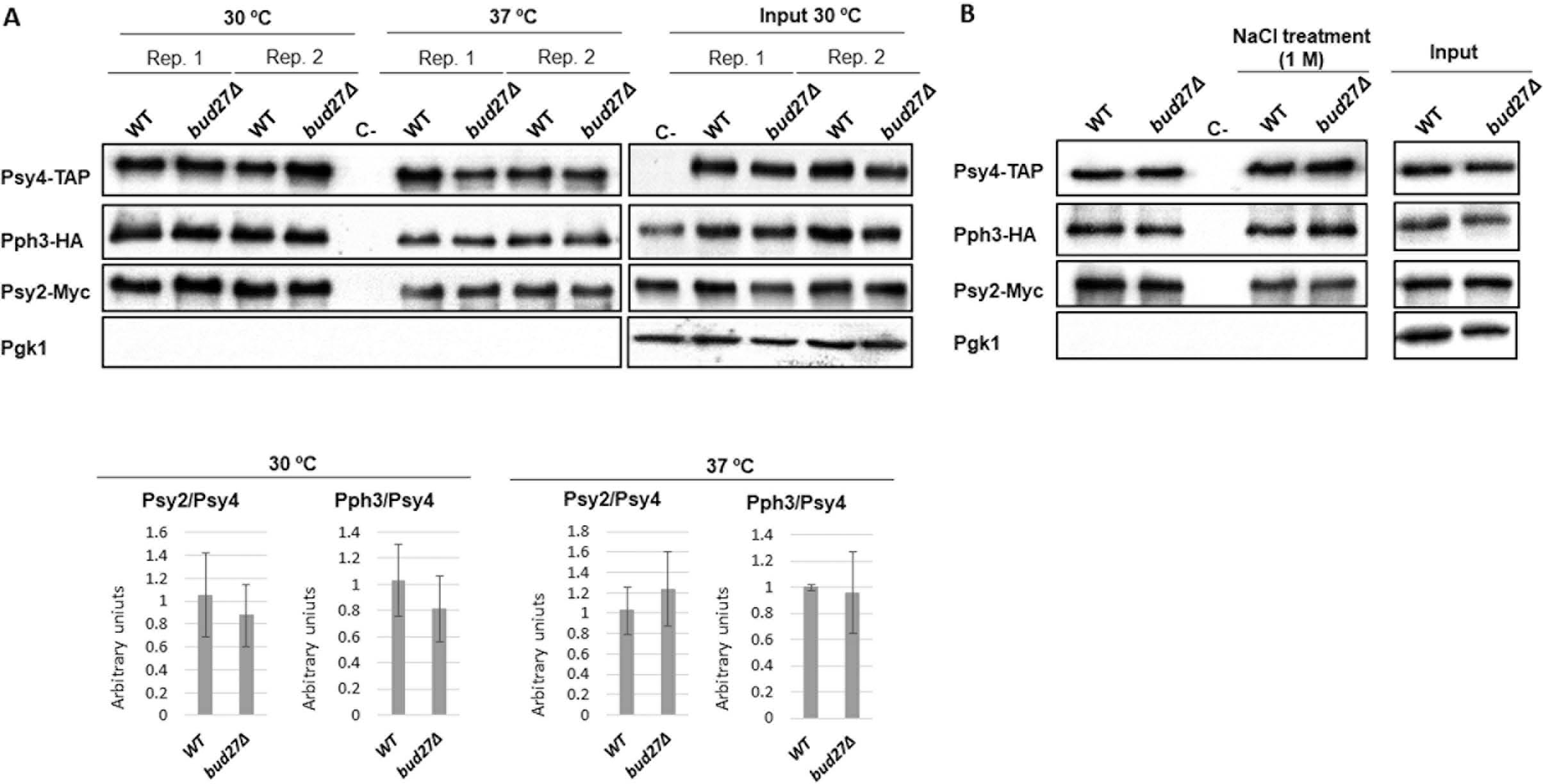
Lack of Bud27 does not appear to affect the stoichiometry or stability of the PP4 complex. A) Upper panel: the whole-cell extracts prepared from the wild-type (WT) *bud27Δ* cells containing a TAP-tagged version of Psy4, a HA-tagged version of Pph3 and a Myc-tagged version of Psy2 were incubated at 30°C or 37°C for 30 minutes and then subjected to pull-down with Dynabeads Pan-Mouse. The obtained immunoprecipitates were analysed by western blotting with antibodies against TAP, HA, Myc or Pgk1. Two independent biological replicates (Rep.) of the experiment are shown. A TAP-untagged strain (C-) was used as a control. Bottom panel. Quantification of the western blottings shown in the upper panel quantified by densitometry. Graphs represent the median and standard deviation of two independent biological replicates. B) The whole-cell extracts prepared from the wild-type (WT) or *bud27Δ* cells containing Psy4-TAP, Pph3-HA and Psy2-Myc were incubated (or not) with increasing NaCl concentrations (250 mM, 500 mM and 1 M) before immunoprecipitation using Dynabeads Pan-Mouse. The obtained immunoprecipitates were analysed by western blotting with antibodies against TAP, HA, Myc, or Pgk1. A TAP-untagged strain (C-) was used as a control.

To analyse the stability of the PP4 complex, protein extracts were incubated at 37°C for 30 minutes before PP4 immunopurification. As shown in Figure 6A (37°C), increasing the temperature did not alter PP4 subunit composition or stoichiometry, which suggests no significant impact of lack of Bud27 on PP4 complex stability. We also analysed the effect of adding salt during PP4 purification by means of three successive washing steps by increasing NaCl concentrations up to 1 M. Once again, no difference in the amounts of the PP4 subunits was observed in the absence of Bud27 (Figure 7B). Altogether, these results suggest that Bud27 may not play a role in PP4 stability and is probably not a co-chaperone of this phosphatase.

### Lack of Bud27 does not alter the cellular localization or phosphorylation of the PP4 complex subunits

The nucleo-cytoplasmic shuttling of the Psy4 subunit of the PP4 complex has been reported to depend on both CDK kinase activity and the cell cycle stage (45,46). Notably, this shuttling is associated with the phosphorylation state of Psy4 (46).

Thus we wondered whether lack of Bud27 could alter the subcellular localization and phosphorylation pattern of PP4 subunits Pph3, Psy2 and Psy4. To do so, we investigated the localization of the PP4 subunits by immunohistochemistry in the wild-type and *bud27Δ* cells expressing Pph3-HA, Psy2-Myc and Psy4-TAP in both the presence and absence of rapamycin.

Our results in Figure 8A revealed that Pph3, Psy2 and Psy4 were mainly nuclear and lack of Bud27 did not seem to alter this localization. Moreover, TOR cascade inhibition did not drastically affect the localization of the PP4 subunits, although rapamycin addition led to slight cytoplasmic Psy4 accumulation in both the wild-type and *bud27Δ* cells.

**Figure 8.**
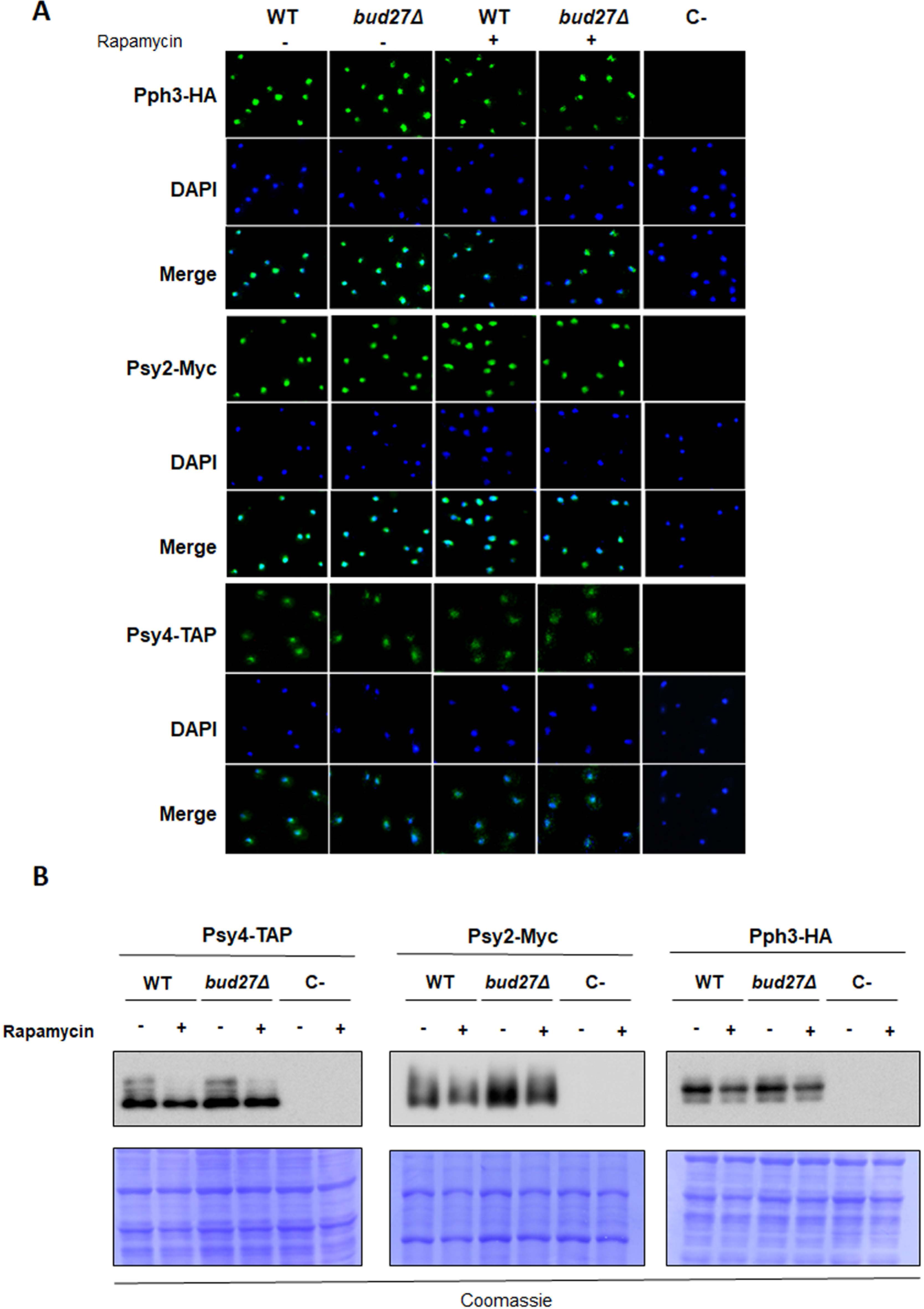
Lack of Bud27 does not affect the localization or phosphorylation of the PP4 subunits. A) Live-cell imaging of Pph3-HA, Psy2-Myc or Psy4-TAP in the wild-type (WT) or *bud27Δ* cells treated with rapamycin for 1 h. An untagged strain was used as a negative control. B) The whole-cell extracts prepared from the cultures used in the experiments shown in panel A were subjected to Phos-Tag SDS-PAGE and analysed by western blotting to determine the phosphorylation state of Psy4, Psy2 and Pph3 upon rapamycin treatment. An untagged strain was used as a negative control. Coomassie blue staining is shown as a loading control.

Furthermore, the analysis of PP4 subunits phosphorylation by Phos-tag SDS-polyacrylamide gel electrophoresis revealed no clear differences between the wild-type and *bud27Δ* cells (Figure 8B). Notably upon rapamycin treatment, a similar decrease in Psy4 phosphorylation was found in both strains.

These results indicate that the defect of PP4 activity on Maf1 in *bud27Δ* cells does not seem to depend on the localization of the PP4 subunits or their phosphorylation pattern.

### PP4 phosphatase activity depends on Bud27

Our data suggest a functional relation between Bud27 and PP4 phosphatase to regulate Maf1 dephosphorylation. We wondered whether the defect in PP4 phosphatase activity was specific to Maf1 or if it could affect other PP4 targets. For this purpose, we investigated Rad53 phosphorylation levels after inducing a single DSB (double strand break) in chromosome III as Rad53 dephosphorylation has been shown to be controlled by PP4 during the repair of these DNA lesions (33).

We analysed Rad53 phosphorylation state by western blotting (Figure 9A upper panel) and the cell cycle by FACS (Figure 9A lower panel) in either the *bud27Δ*, *pph3Δ* and *bud27Δ pph3Δ* mutant cells or the isogenic wild-type strain after inducing a single DSB by adding galactose to the culture as previously described (33). As expected (33), Rad53 was first hyperphosphorylated with a maximum observed 6 h after DSB generation, and was then progressively dephosphorylated to reach its basal phosphorylation state after 11 h in the wild-type cells. Lack of Bud27 led to a prolonged phosphorylation Rad53 state with a maximum detected 8 h after DSB induction, and dephosphorylation kinetics clearly slowed down. Accordingly, the *bud27Δ* cells blocked the G2/M DNA damage checkpoint and re-entered the cell cycle with delayed kinetics compared to the control strain (Figure 9A, lower panel). Similar defects in Rad53 dephosphorylation and cell cycle re-entry were observed for the *pph3Δ* mutant, a finding that corroborates what has been previously reported (33). However, the defects in Rad53 dephosphorylation were exacerbated in the *pph3Δ* cells compared to *bud27Δ* (Figure 9A). This scenario suggests that although PP4 activity was compromised in the absence of Bud27, the *bud27Δ* cells still retained some phosphatase complex activity. Interestingly, the double *pph3Δ bud27Δ* mutant displayed similar Rad53 phosphorylation and cell cycle re-entry profiles to the single *pph3Δ* mutant, which indicates that the defects observed in the *bud27Δ* cells were probably only due to decreased PP4 activity.

**Figure 9.**
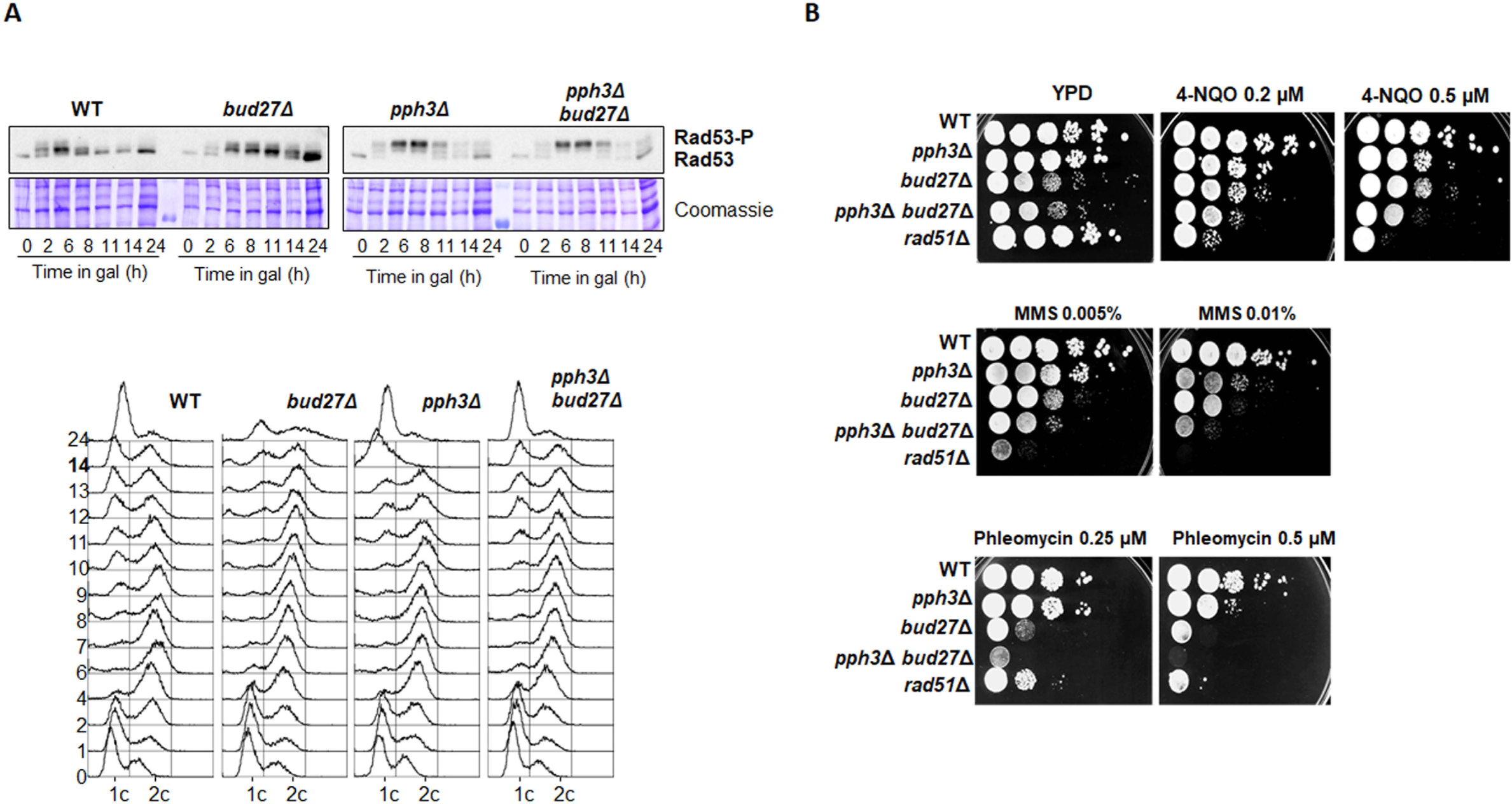
Bud27 influences PP4 activity under DNA-damaging conditions. A) Top: the cultures of the YMV80 (WT) strain or its *bud27Δ*, *pph3Δ* and *pph3Δ bud27Δ* derivative were supplemented with galactose to induce the expression of the HO nuclease to generate a double strand break in chromosome III. Samples were collected at the indicated time points and Rad53 phosphorylation levels were analysed by western blotting. Coomassie blue staining is shown as a loading control. Bottom: the FACS profiles of the samples collected from the cultures used in panel A were processed to determine DNA content throughout the experiment. B) Genetic interactions between *BUD27* and *PPH3*. Serial dilutions of the indicated cultures were spotted onto YPD plates containing, or not, the indicated concentration of DNA-damaging agents 4-nitroquinoline-1-oxide (4-NQO), methyl methanesulphonate (MMS) or phleomycin. The *rad51Δ* strain was used as an internal control.

These results show that Bud27 is necessary to maintain steady-state PP4 activity, and to not only promote Maf1 dephosphorylation, but to also contribute to the dephosphorylation of other proteins like Rad53.

As *pph3Δ* mutant cells are sensitive to different DNA-damaging agents (33), we analysed the effect of some genotoxic drugs on the growth of cells lacking Bud27 or/and Pph3 compared to the isogenic wild-type strain using *rad51Δ* as a control. The *bud27Δ* mutant showed slightly more sensitivity than the *pph3Δ* mutant to UV-mimetic agent 4-nitroquinoline-1-oxide (4-NQO), alkylating agent methyl methanesulphonate (MMS) and radiomimetic drug phleomycin (Figure 9B). Furthermore, the fact that the lack of both Bud27 and Pph3 aggravated cell growth defects in the presence of these genotoxic drugs, indicates that apart from its role in influencing PP4 activity, Bud27 might play other roles in responses to bulky DNA damage agents.

Taken together, these results indicate that Bud27 interferes with PP4 activity in response to a single DSB lesion in DNA. In addition, lack of Bud27 has a stronger effect than lack of PP4 when exposing cells to bulky DNA damage agents, which suggests that Bud27 may participate in the maintenance of other DNA damage-related proteins, possibly through its role as a prefoldin protein.

## Discussion

Bud27 is a prefoldin-like that interacts with the three RNA pols through their shared subunit Rpb5 and plays roles in transcription (11,15). Bud27 modulates RNA pol III transcription through either its participation in the TOR signalling pathway or its association with the RSC remodelling complex, but the molecular mechanisms of its function in RNA pol III transcriptional repression remain largely unknown (7,13,15,16).

In this work, we demonstrate that Bud27 is required to regulate the dephosphorylation and nuclear localization of Maf1. We show that Bud27 associates with Maf1 phosphatase PP4, which suggests a functional role of Bud27-PP4 interaction in the Maf1 phosphorylation state. Bud27 also seems necessary for the interaction between PP4 and Maf1, which implies its likely influence on the repression of RNA pol III transcription.

Maf1 dephosphorylation is similarly altered by either lack of Bud27 or PP4 inactivation upon TOR pathway repression or nutrient deprivation (36). Furthermore, Maf1 dephosphorylation defects are not additive in the double *bud27Δ pph3Δ* mutant, which means that both Bud27 and PP4 act in the same biological process. Our data point out the concerted action of Bud27 and PP4 in Maf1 dephosphorylation that could occur *via* the interaction between these proteins, and could reinforce the role of PP4 as the main Maf1 phosphatase (36). In line with previously proposed findings (36), this interaction likely occurs in the nucleus and with Bud27 (3,4) and its URI orthologues (1,2) as nuclear proteins. Our data confirm that the Psy2 and Psy4 subunits of PP4 are mainly nuclear proteins (36,45), and they reveal that it is also the case of Pph3 in both the wild-type and *bud27Δ* cells. Interestingly, we show that blocking the TOR pathway by rapamycin addition leads to the partial relocalization of Psy4 in the cytoplasm, which indicates that Psy4 could exist in the cell independently of the core Pph3-Psy2-Psy4, which agrees with previously reported data (36). Our results also demonstrate a novel interaction of Bud27 with a component of the TOR signalling pathway, and they shed light on the role of this prefoldin-like, and likely its human orthologue URI, in this cascade (13,16).

The interaction between Bud27 and PP4 phosphatase to regulate Maf1 dephosphorylation suggests that similar mechanisms may act for other phosphatase-dependent processes, which is the case of human URI and PP2A phosphatase during KAP1 phosphorylation regulation (14). Our data are also indicative that Bud27 is necessary for the interaction between PP4 and Maf1 before its dephosphorylation (Figure 10). It is worth noting that the PP4-Maf1 interaction has also been previously reported (36). Lack of Bud27 decreases the PP4-Maf1 association, which probably accounts for the decrease in Maf1 dephosphorylation observed by us or others upon PP4 inactivation (19,36). Given that URI is phosphorylated (1), and considering that Bud27 has putative phosphorylation sites, we speculate that Bud27 might play a role in the posttranslational modification of PP4 (phosphorylation) that is necessary for Maf1 dephosphorylation. However, if this were indeed the case, the modification may be subtle because no differences were detected in Phos-Tag gel analyses. Furthermore, in line with the role of Bud27 as a co-chaperone (1), it is tempting to speculate that Bud27 could mediate the necessary folding of PP4 once both are associated by ensuring its interaction with Maf1 and subsequent Maf1 dephosphorylation. This mechanism would fall in line with that proposed for URI/PP2A/KAP1 (14). However, the fact that our data show no differences in PP4 stoichiometry and suggest no major defect in PP4 stability casts doubt about this possible mechanism. In addition, although we were unable to detect any interaction between Bud27 and Maf1, we cannot rule out a tripartite interaction that would involve Bud27, PP4 and Maf1, which may be necessary for Maf1 dephosphorylation.

**Figure 10.**
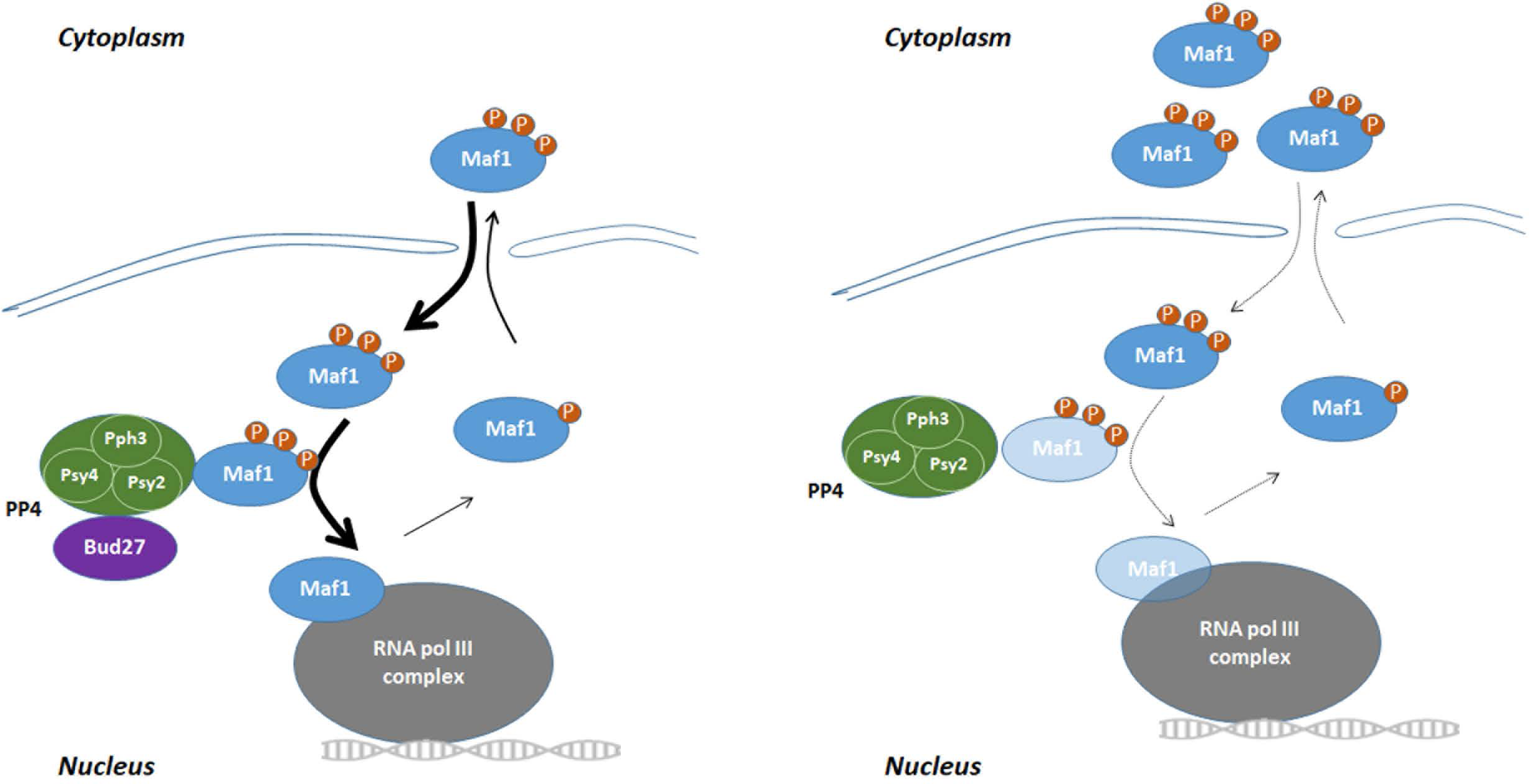
Model for Bud27-PP4-dependent Maf1 dephosphorylation. Bud27 interacts with PP4 to allow proper Maf1 dephosphorylation and nuclear entry that, in turn, mediates correct RNA pol III transcriptional repression (left image). In the absence of Bud27, both Maf1 nuclear entry and the interaction between Maf1 and PP4 decreases affect Maf1 dephosphorylation and RNA pol III repression (right image).

In line with the cytoplasmic-nuclear shuttling of Maf1 that depends on its dephosphorylation (19,24,36), lack of Bud27 decreases Maf1 nuclear accumulation. Furthermore, and as expected, in the *bud27Δ* mutant the rapid RNA pol III repression seemed to be affected in a similar way to that in other situations which alter Maf1 dephosphorylation (19,24,36,47). However, it is noteworthy that lack of Bud27 had additional effects on RNA pol III transcriptional machinery by affecting RNA synthesis and likely processing (Gutiérrez-Santiago *et al*, in preparation). This could potentially interfere with the rapid repression that occurred after Maf1 dephosphorylation. Furthermore, this additional defect on RNA pol III in the *bud27Δ* mutant cells made the tRNA repression analysis more complex. We speculate that the association between Bud27 and PP4 in the nucleus could account for the rapid repression of RNA pol III that occurred under stress. However, the effect of lack of Bud27 on tRNA repression could vary depending on the analysed tRNA, and in agreement with the differences proposed for the synthesis and maturation of different types of tRNAs in yeast and human cells (48-52). Notably, and in line with Bud27-PP4 acting on RNA pol III regulation through transcriptional repression, human URI influences transcription repression in cancer prostate cells *via* PP2A phosphatase (14).

Our data also suggest a more general effect of Bud27 on PP4 activity because not only Maf1, but also Rad53 dephosphorylation, are altered upon DSB lesion in DNA (33). Accordingly, Bud27 may be necessary for PP4 activity to act on several targets, which implies similar mechanisms to those proposed for Maf1 dephosphorylation (posttranslational modification or folding). However, we cannot rule out an additional role for Bud27 in DNA damage in line with previous data showing a relation between Bud27 and URI with DNA damage, DNA stability or DNA integrity (47,53-55).

## Acknowledgments

We thank Dr. Jesús de la Cruz, Dr. Olga Rodríguez-Galán and the “Servicios Centrales de Apoyo a la Investigación (SCAI)” of the University of Jaén for technical support.

