## Supplementary Material for "Maf1 phosphorylation is regulated through the action of prefoldin-like Bud27 on PP4 phosphatase in *Saccharomyces cerevisiae*"

### Table S1. *S. cerevisiae* strains

| Strain | Genotype | Origin |
|---|---|---|
| AC1327 | *HOΔ hml::ADE1 mata::hisG hmr::ADE1 his4::NAT-leu (Xho- to Asp718) leu2::HOcs ade3::GAL::HO ade1 lys5 ura3-52 trp1::hisG pph3Δ::HPH* | (1) |
| AC1822 | *HOΔ hml::ADE1 mata::hisG hmr::ADE1 his4::NAT-leu-(Xho- to Asp718) leu2::HOcs ade3::GAL::HO ade1 lys5 ura3-52 trp1::hisG* | (1) |
| AC1468 | *HOΔ hml::ADE1 mata::hisG hmr::ADE1 his4::NAT-leu-(Xho- to Asp718) leu2::HOcs ade3::GAL::HO ade1 lys5 ura3-52 trp1::hisG rad51Δ::URA* | (1) |
| BY4741 | *MATa his3-Δ1 leu2-Δ0 met15-Δ0 ura3-Δ0* | Euroscarf |
| BY4742 | *MATα his3-Δ1 leu2-Δ0 lys2-Δ0 ura3-Δ0* | Euroscarf |
| MAF1-GFP | *MATa his3-Δ1 leu2-Δ0 met15-Δ0 ura3-Δ0 MAF1::GFP::HIS3MX6* | Gift from M. Boguta |
| YFN683 | *MATa his3-Δ1 leu2-Δ0 met15-Δ0 ura3-Δ0 YFL023W::natNT2 (bud27::natNT2)* | This work |
| YFN684 | *MATα his3-Δ1 leu2-Δ0 lys2-Δ0 ura3-Δ0 YJL164C::kanMX4 (tpk1::kanMX4) YFL023W::natNT2 (bud27::natNT2)* | This work |
| YFN685 | *MATα his3-Δ1 leu2-Δ0 lys2-Δ0 ura3-Δ0 YPL203W::kanMX4 (tpk2::kanMX4) YFL023W::natNT2 (bud27::natNT2)* | This work |
| YFN686 | *MATα his3-Δ1 leu2-Δ0 lys2-Δ0 ura3-Δ0 YKL166C::kanMX4 (tpk3::kanMX4) YFL023W::natNT2 (bud27::natNT2)* | This work |
| YFN687 | *MATα his3-Δ1 leu2-Δ0 lys2-Δ0 ura3-Δ0 YHR205W::kanMX4 (sch9::kanMX4) YFL023W::natNT2 (bud27::natNT2)* | This work |
| YFN688 | *MATα his3-Δ1 leu2-Δ0 lys2-Δ0 ura3-Δ0 YAL016W::kanMX4 (tpd3::kanMX4) YFL023W::natNT2 (bud27::natNT2)* | This work |
| YFN689 | *MATα his3-Δ1 leu2-Δ0 lys2-Δ0 ura3-Δ0 YIL035C::kanMX4 (cka1::kanMX4) YFL023W::natNT2 (bud27::natNT2)* | This work |
| YFN690 | *MATα his3-Δ1 leu2-Δ0 lys2-Δ0 ura3-Δ0 YPL180W::kanMX4 (tco89::kanMX4) YFL023W::natNT2 (bud27::natNT2)* | This work |
| YFN691 | *MATα his3-Δ1 leu2-Δ0 lys2-Δ0 ura3-Δ0 YIL035C::kanMX4 (pph3::kanMX4) YFL023W::natNT2 (bud27::natNT2)* | This work |

| | | |
|---|---|---|
| YFN734 | *MATa his3- Δ1 leu2- Δ0 met15- Δ0 ura3- Δ0 MAF1::GFP::HIS3MX6 YFL023W::natNT2 (bud27::natNT2)* | This work |
| YFN778 | *MATα his3-Δ1 leu2-Δ0 lys2-Δ0 ura3-Δ0 YDR007w::kanMX4 (TRP1::kanMX4) YFL023W::natNT2 (bud27::natNT2)* | This work |
| YFN787 | *MATa his3-Δ1 leu2-Δ0 met15-Δ0 ura3-Δ0 PPH3::3xHA::HIS3MX6 PSY2::13xMyc::kanMX6* | This work |
| YFN788 | *MATa his3-Δ1 leu2-Δ0 met15-Δ0 ura3-Δ0 PPH3::3xHA::HIS3MX6 PSY2::13xMyc::kanMX6 YFL023W::natNT2 (bud27::natNT2)* | This work |
| YFN800 | *MATa his3-Δ1 leu2-Δ0 met15-Δ0 ura3-Δ0 PPH3::3xHA::HIS3MX6 PSY2::13xMyc::kanMX6 PSY4::TAP::URA3* | This work |
| YFN801 | *MATa his3-Δ1 leu2-Δ0 met15-Δ0 ura3-Δ0 PPH3::3xHA::HIS3MX6 PSY2::13xMyc::kanMX6 PSY4::TAP::URA3 YFL023W::natNT2 (bud27::natNT2)* | This work |
| YFN802 | *HOΔ hml::ADE1 mata::hisG hmr::ADE1 his4::NAT-leu-(Xho- to Asp718) leu2::HOcs ade3::GAL::HO ade1 lys5 ura3-52 trp1::hisG YFL023W::kanMX4 (bud27::kanMX4)* | This work |
| YFN803 | *HOΔ hml::ADE1 mata::hisG hmr::ADE1 his4::NAT-leu-(Xho- to Asp718) leu2::HOcs ade3::GAL::HO ade1 lys5 ura3-52 trp1::hisG pph3Δ::HPH YFL023W::kanMX4 (bud27::kanMX4)* | This work |
| YFN815 | *MATα his3-Δ1 leu2-Δ0 lys2-Δ0 ura3-Δ0 YDR007w::kanMX4 (TRP1::kanMX4) PPH3-MYC::TRP1* | This work |
| YFN831 | *MATα his3-Δ1 leu2-Δ0 lys2-Δ0 ura3-Δ0 YDR007w::kanMX4 (TRP1::kanMX4) PPH3-MYC::TRP1 BUD27-LytA::HIS3* | This work |
| Y11089 | *MATα his3-Δ1 leu2-Δ0 lys2-Δ0 ura3-Δ0 YPL203W::kanMX4 (tpk2::kanMX4)* | Euroscarf |
| Y11261 | *MATα his3-Δ1 leu2-Δ0 lys2-Δ0 ura3-Δ0 YJL164C::kanMX4 (tpk1::kanMX4)* | Euroscarf |
| Y11428 | *MATα his3-Δ1 leu2-Δ0 lys2-Δ0 ura3-Δ0 YIL035C::kanMX4 (cka1::kanMX4)* | Euroscarf |
| Y12072 | *MATα his3-Δ1 leu2-Δ0 lys2-Δ0 ura3-Δ0 YPL180W::kanMX4 (tco89:: kanMX4)* | Euroscarf |
| Y14010 | *MATα his3-Δ1 leu2-Δ0 lys2-Δ0 ura3-Δ0 YIL035C::kanMX4 (pph3::kanMX4)* | Euroscarf |
| Y15016 | *MATα his3-Δ1 leu2-Δ0 lys2-Δ0 ura3-Δ0 YKL166C::kanMX4 (tpk3::kanMX4)* | Euroscarf |
| Y16866 | *MATα his3-Δ1 leu2-Δ0 lys2-Δ0 ura3-Δ0 YAL016W::kanMX4 (tpd3::kanMX4)* | Euroscarf |

| | | |
|---|---|---|
| Y17202 | *MATα his3-Δ1 leu2-Δ0 lys2-Δ0 ura3-Δ0 YDR007W::kanMX4 (TRP1::kanMX4)* | Euroscarf |
| Y17797 | *MATα his3-Δ1 leu2-Δ0 lys2-Δ0 ura3-Δ0 YHR205W::kanMX4 (sch9::kanMX4)* | Euroscarf |
| YSS2 | *MATα ade2-101 lys2-801 ura3-52 trp1-Δ63 his3-Δ200 RPO31::13Myc::kanMX4* | (2) |
| YFNOL1 | *MATα ade2-101 lys2-801 ura3-52 trp1-Δ63 his3-Δ200 RPO31::13Myc::kanMX4 YFL023W::HIS3 (bud27::HIS3)* | (2) |

### Table S2. Plasmids

| **Name** | **Yeast Marker** | **Origin** |
|---|---|---|
| pCM185 | ORI (CEN) *TRP1* | (3) |
| pCM185-*PPH3*-TAP | ORI (CEN) *TRP1* | This work |
| pFL44L-*RPB5* | ORI (2µm) *URA3* | (4) |
| pFL44L-*MAF1* | ORI (2µm) *URA3* | Gift from M. Boguta |
| pFL44L | ORI (2µm) *URA3* | (5) |

### Table S3. Oligonucleotides used in this study for RT-qPCR

| Gene (or DNA region) | Primer | Sequence |
|---|---|---|
| ***18S rDNA*** | 18S-501 | CATGGCCGTTCTTAGTTGGT |
| | 18S-301 | ATTGCCTCAAACTTCCATCG |
| **tRNALeu3** | tRNALeu3-302 (Used for RT) | GAGATTCGAACTCTTGCATCTT |
| | pretRNALeu3-502 | GGCGCCTGATTCAAGAAATA |
| | tRNALeu3-301 | TTCGAACTCTTGCATCTTACG |
| **tRNATrp** | tRNATrp-302 (Used for RT) | TGAAACGGACAGGAATTGAACC |
| | pretRNATrp-501 | CGACTCCAATTAAATCTTGGAAA |
| | tRNATrp-301 | GGAATTGAACCTGCAACCCT |

### Table S4. Oligonucleotides used in this study for northern blot

| Probe | Sequence |
|---|---|
| Probe e (5.8S) | TTTCGCTGCGTTCTTCATC |
| tRNALeu(CAA)A-(Sup56) | CCTTAGACCGCTCGGCCAAA |
| tRNAIle(AAU)I | TGCTCGAGGTGGGGTTGAACCCACGACGG |

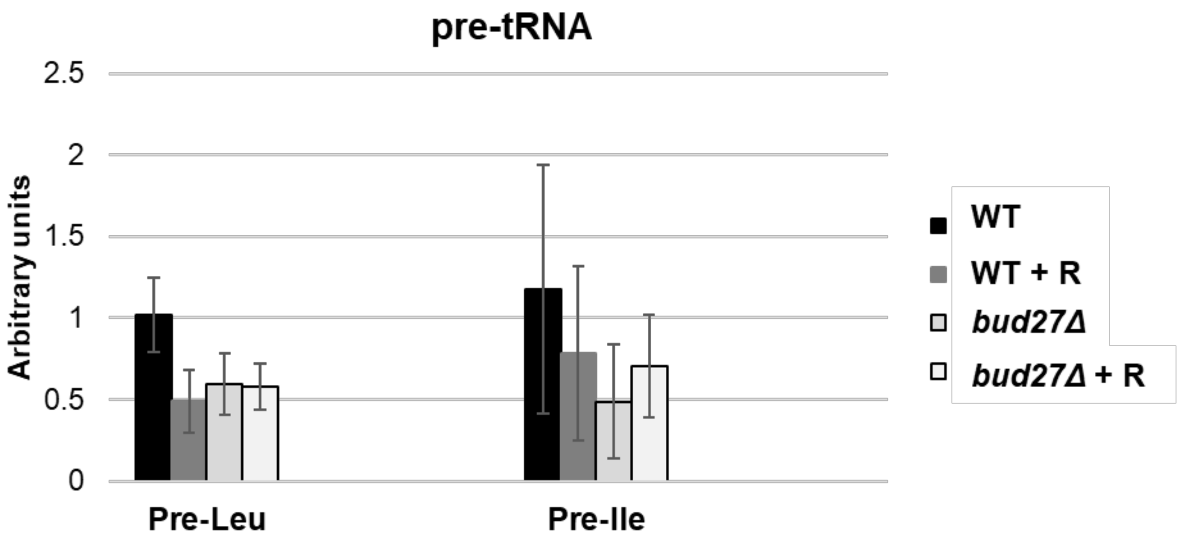

Figure S1

**Figure S1. Lack of Bud27 affects RNA pol III repression.** Northern blot to analyse neosynthesis of tRNALeu3 (pre-leu) and tRNAIle (pre-Ile) from wild-type or bud27Δ cells grown to the log phase and then treated (or not) with rapamycin for 1 hour. rRNA 5.8S was used as a normalizer. Experiments corresponded to two independent biological replicates. Data are the median and standard deviation. +R: Rapamycin treatment
